# Biogeographic Distribution of Five Antarctic Cyanobacteria Using Large-Scale k-mer Searching with sourmash branchwater

**DOI:** 10.1101/2022.10.27.514113

**Authors:** Jessica Lumian, Dawn Sumner, Christen Grettenberger, Anne D. Jungblut, Luiz Irber, N. Tessa Pierce-Ward, C. Titus Brown

## Abstract

Cyanobacteria form diverse communities and are important primary producers in Antarctic freshwater environments, but their geographic distribution patterns in Antarctica and globally are still unresolved. There are however few genomes of cultured cyanobacteria from Antarctica available and therefore metagenome-assembled genomes (MAGs) from Antarctic cyanobacteria microbial mats provide an opportunity to explore distribution of uncultured taxa. These MAGs also allow comparison with metagenomes of cyanobacteria enriched communities from a range of habitats, geographic locations, and climates. However, most MAGs do not contain 16S rRNA gene sequences, making a 16S rRNA gene-based biogeography comparison difficult. An alternative technique is to use large-scale k-mer searching to find genomes of interest in public metagenomes.

This paper presents the results of k-mer based searches for 5 Antarctic cyanobacteria MAGs from Lakes Fryxell and Lake Vanda, assigned the names *Phormidium pseudopriestleyi*, a *Microcoleus*, a *Leptolyngbya*, a *Pseudanabaena*, and a *Neosynechococcus* (Lumian et al., 2021, Lumian et al., 2022, in prep.) in 498,942 unassembled metagenomes from the National Center for Biotechnology Information (NCBI) Sequence Read Archive (SRA). The *Microcoleus* MAG was found in a wide variety of environments, *P. pseudopriestleyi* was found in environments with challenging conditions, the *Neosynechococcus* was only found in Antarctica, and the *Leptolyngbya* and *Pseudanabaena* MAGs were found in Antarctic and other cold environments. The findings based on metagenome matches and global comparisons suggest that these Antarctic cyanobacteria have distinct distribution patterns ranging from locally restricted to global distribution across the cold biosphere and other climatic zones.

## INTRODUCTION

Cyanobacteria are a diverse group of oxygenic photosynthetic gram-negative bacteria that are prevalent in a wide range of environments. In polar environments, cyanobacteria play an important part in shaping local ecology because of their role as primary producers underlying habitats such as benthic biofilms (Stal, 2007; Quesada & Vincent, 2012; Chrismas et al., 2016). Cyanobacteria that thrive in Antarctica face many challenges including variable light availability, cold temperatures, and freeze-drying conditions. To withstand these conditions, cyanobacteria may have tolerance mechanisms encoded in their genomes (Chrismas et al., 2015, 2016). However, the presence of tolerance genes in their genomes may make it more difficult for polar cyanobacteria to compete with other cyanobacteria in non-polar environments. Consequently, some polar cyanobacteria may only occur in polar environments, while others may also be present in environments that share similar conditions to the stresses they face in Antarctica, such as cold temperatures or light stress (Jungblut et al., 2016; Chrismas, Williamson, et al., 2018b; Lumian et al., 2021).

Currently, polar cyanobacteria are underrepresented in genomic databases, despite the important role they play in primary productivity including in perennially ice-covered lakes in the McMurdo Dry Valleys in Antarctica. Due to a lack of grazers and limited water mixing, vast microbial mats in the Lakes Vanda and Fryxell prosper and sustain complex geochemical gradients (Jungblut et al., 2016; Sumner et al., 2016; Lumian et al., 2021). These geochemical gradients structure competition within communities, which are also dealing with challenging environmental conditions, such as highly seasonal light, nutrient, limitation, and in part of Lake Fryxell, sulfidic water (Lumian et al., 2021; Jungblut et al. 2016; Dillon et al. 2020).

The question of why Antarctic cyanobacteria can survive in challenging conditions and what other environments they grow in can be addressed by biogeography studies (Whitaker et al., 2003; Martiny et al., 2006; Fierer, 2008; Green et al. 2008). Previous 16S rRNA gene surveys based on amplicon sequencing provided support for the longstanding theory that microbes have unlimited dispersal and community distribution is selected by the environment (Baas-Becking, 1934; Jungblut et al. 2010; Harding et al 2011). However, studies from other environments and climatic zones have shown that 16S rRNA gene and single gene markers might not provide sufficient resolution to resolve genotypes and populations.

Most biogeography studies on polar microbiomes and cyanobacteria to date are based on 16S rRNA gene amplicon sequencing in the context of local environmental conditions of sampling sites or pole-to-pole comparisons using clone library surveys and high throughput sequencing approaches (Taton et al., 2006; Namsaraev et al., 2010; Jungblut et al. 2011; Bahl et al., 2011; Moreira et al., 2013; Harke et al., 2016; Kleinteich et al. 2017; Ribeiro et al., 2018). Although 16S rRNA gene sequences are computationally easier to compare to each other, there are limitations to 16S rRNA gene-based biogeography studies. The 16S rRNA gene is conserved and therefore likely leads to an under estimation of genotype level richness. Furthermore, the short read length of high throughput sequencing only allows the coverage of a few variable regions which further reduces phylogenetic resolution. While recent genomic work has provided advances in biogeography of polar microbes (Chrismas et al. 2015), the 16S rRNA gene sequence may not assemble and bin well from metagenomes, which can prohibit MAGs from being incorporated into 16S rRNA gene-based biogeographical distributions.

An alternative to 16S rRNA gene-based biogeography is to apply comparative genomic approaches, but this is computationally more complicated because of the size and scale of metagenome datasets. One option is to use an alignment-based approach in which the reads are aligned to reference genomes, which has been done for large-scale viral genome discovery with Serratus (Edgar et al., 2022). Another option is to apply large-scale k-mer matching to unassembled metagenomes, which is possible with sourmash branchwater (Irber et al., 2022; Brown & Irber, 2016; Irber, 2020a, 2020b; Brown, 2021). These techniques open the possibility of using metagenomic data for biogeography studies by searching all publicly available metagenomes on the National Center for Biotechnology Information (NCBI) Sequence Read Archive (SRA) (Leinonen et al., 2011) for Antarctic MAGs of interest. In this paper, branchwater was used to search 498,942 unassembled metagenomes from the NCBI SRA for the presence of five Antarctic cyanobacteria MAGs that lack the 16S rRNA gene. These findings provide new insights based on comparative genomic analyses into the distribution patterns of cyanobacteria in cold biospheres.

## MATERIALS AND METHODS

### Selection of Antarctic Cyanobacteria of Interest

*Phormidium pseudopriestleyi* is a well characterized cyanobacteria in Lake Fryxell, Antarctica (Lumian et al., 2021). Lake Fryxell is a perennially ice-covered lake located at 77.36° S, 162.6° E in the McMurdo Dry Valleys. The base of the lake is covered with microbial mats, with *P. pseudopriestleyi* dominating the mats at 9.8 m in depth, where light levels are low (1-2 µmol photons m^−2^ s^−1^) and sulfide is present in the water column (0.091 mg L^−1^). *P. pseudopriestleyi* performs oxygenic photosynthesis in the presence of hydrogen sulfide, even though sulfide inhibits oxygenic photosynthesis (Sumner et al., 2015; Lumian et al., 2021). Lake conditions and sampling have been described in Jungblut et al. (2016), Dillon et al. (2020), and Lumian et al. (2021).

The *Neosynechococcus, Leptolyngbya, Microcoleus*, and *Pseudanabaena* MAGs are from microbial mats located in Lake Vanda, McMurdo Dry Valleys. Lake Vanda is also a perennially ice-covered lake and is located at 77.53° S, 161.58° E. Microbial mats in Lake Vanda contain pinnacles that range from millimeters to centimeters tall. Unlike Lake Fryxell, there is no sulfide where we sampled, and it is better illuminated at the sampled location than Lake Fryxell, though samples from the inside of pinnacles receive little light (Sumner et al., 2016). Sampling methods and lake conditions have previously been described in Sumner et al. (2016)

### Bioinformatic Processing and Assembly of Antarctic Cyanobacteria Reference MAGs

The methods to obtain the *P. pseudopriestleyi* MAG has been previously described in Lumian et al. (2021). Briefly, the *P. pseudopriestleyi* MAG was obtained from a microbial mat sample sequenced on an Illumina HiSeq 2500 PE250 platform and a laboratory culture was sequenced on an Illumina 2000 PE100 platform. The microbial mat sample was quality filtered, and forward and reverse reads were joined using PEAR v0.9.6 (Zhang et al., 2014). For the isolated strain, trimmomatic v0.36 (Bolger et al., 2014) was used to trim sequencing adapters, and the interleave-reads.py script in khmer v2.1.2 (Crusoe et al., 2015) was used to interleave the reads. Both samples were assembled separately and together as a co-assembly by MEGAHIT v1.1.2 (Li et al., 2015) and mapped with bwa v2.3 (Li, 2013) and samtools v1.9 (Li et al., 2009). A single cyanobacteria bin was obtained using the CONCOCT binning algorithm in anvi’o and identified using CheckM (Eren et al., 2015; Delmont & Eren, 2018; Parks et al., 2015). The *P. pseudopriestleyi* bin was refined with spacegraphcats to extract additional content from the metagenomes with a k-mer size of 21 and a radius of 1 (Brown et al., 2020).

Methods to obtain the *Microcoleus, Pseudanabaena, Leptolyngyba*, and *Neosynechococcus* MAGs from Lake Vanda are described in Lumian et al. 2022 (unpublished). Filtered and quality controlled raw data was retrieved from the NCBI Sequence Read Archive under the accession numbers SRR6448204 - SRR6448219 and SRR 6831528.. MEGAHIT v1.9.6 was used to assemble metagenomes with a minimum contig length of 1500 bp and a paired end setting. Bowtie2 v1.2.2 and samtools v1.7 were used to map reads back to the assembly. A depth file was generated using jgi_summarize_bam_contig_depths from MetaBAT v2.12.1 (Kang et al., 2015), which was also used to generate bins with a minimum contig length of 2500 bp. The completeness and contamination of the bins were calculated with CheckM v1.0.7 (Parks, D.H., et al., 2014). Bins that were contained within the phylum Cyanobacteria in the phylogenetic tree generated by CheckM were retained for further analysis. 139 single copy marker genes (Campbell et al., 2013) were collected using the anvi-run-hmms command in anvi’o v6.2 (Eren et al., 2021) and a phylogenetic tree was constructed using the anvi-gen-phylogenomic-tree command. Genome similarity between bins was computed using the anvi-compute-genome-similarity command. Bins were grouped into taxa if they shared more than 98% average nucleotide similarity and were phylogenetically cohesive. When a taxon was found in multiple metagenomes, the most complete bin with the lowest level of contamination for that taxon was selected for additional analysis. <<Could say something like the bin selected for each taxon will be referred to as the MAG for that taxon -- something to explain why we jump from the phrase “bin” to “MAG”>> Taxa were classified using GTDB-tk v.2.1.0 (Chaumeil et al., 2020). MAGs for each taxon are being deposited in the NCBI sequence read archive under and will receive accession numbers.

### Sourmash branchwater Software with Large-Scale k-mer Searching for Comparative Metagenomic Analysis

The branchwater software used large-scale k-mer searching to search all metagenomes in the NCBI SRA as of September 2020 for matches with genomes of interest (Brown & Irber, 2016; Pierce et al., 2019). Signature files of the genomes of interest were generated using sourmash v3.5.0 (Brown & Irber, 2016) with k-mer sizes of 21, 31, 51, the scaled parameter set to 1000, and abundance tracking. This generated a unique signature file specific to each of the five Antarctic MAGs. These signature files were searched against signature files previously generated for all 498,942 publicly available unassembled metagenome sets on the SRA as of September, 2020 using exact k-mer matching. Results are organized by containment, which is the proportion of the query MAG k-mers found in the metagenome. The use of k = 31 as a k-mer size enables detection of matches to ∼91% ANI at 5% containment and ∼97% ANI at 30% containment (Hera et al., 2022). The size of the Antarctic query MAGs ranged from 2.7 Mbp – 6.07 Mbp, so a match with containment value of 5% implies 135,000 – 303,500 matching k-mers with k = 31 and 4,185,000 – 9,408,500 matching base pairs, which indicates significant shared genomic material between MAGs and metagenome matches.

Validation of k-mer results from branchwater was done by mapping the Antarctic MAGs back to the metagenomes from the SRA using minimap2 v2.24 in genome-grist v0.8.4 (Li, 2018; Irber et al., 2022). Environmental metadata for the top hits of all MAGs with hits above 5% were recorded, with the exception of the *Microcoleus*, which had over 1,000 matches above that threshold (Tables 2 – 3). Unassembled metagenomes from geographically distinct environments were assembled with MEGAHIT v1.9.6, mapped with bowtie2 v1.2.2 and samtools v1.7, and binned with MetaBAT v.2.12.1. However, none of the assemblies were high enough quality to yield bins (Table 4). The code from this project is available at: https://github.com/dib-lab/2022-pipeline-antarctic-biogeography.

## RESULTS

The five polar cyanobacteria MAGs used as search queries were found in a variety of non-polar metagenomic data sets in a range of environmental conditions (Table 1, Figure 1). The metagenome data sets with >5% containment of the MAGs described in Tables 1 – 3. Information about additional environments where the *Microcoleus* MAG was found with over 20% containment is displayed in Table 3. Validation mapping numbers are available in Table 5. The SRA accession numbers of additional hits are available in Supplementary Tables S1 – S5.

**Table 1.** Summary of branchwater hits

|  | *Microcoleus* | *Phormidium pseudopriestleyi* | *Pseudanabaena* | *Neosynechococcus* | *Leptolyngbya* |
| --- | --- | --- | --- | --- | --- |
| # Hits >75% Containment | 6 | 30 | 6 | 3 | 5 |
| # Hits >50% Containment | 12 | 33 | 6 | 5 | 5 |
| # Hits >25% Containment | 119 | 38 | 10 | 6 | 6 |
| # Hits >5% Containment | 1,121 | 131 | 24 | 16 | 22 |
| Total Hits | 6,184 | 2,739 | 3,769 | 2,796 | 2,999 |
| # Geographically Distinct Locations >25% Containment | 27 | 3 | 3 | 1 | 1 |

**Table 2.**
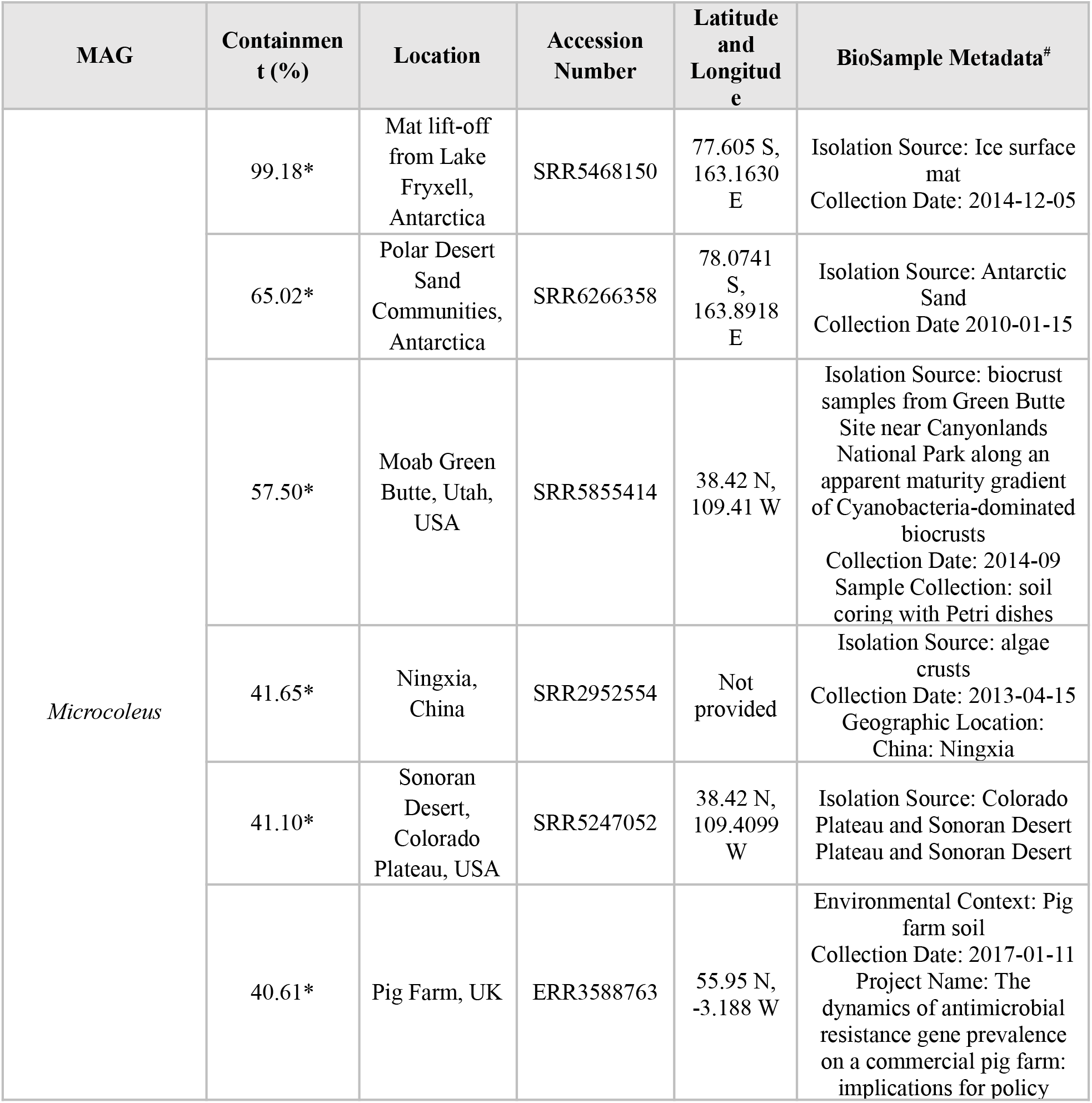
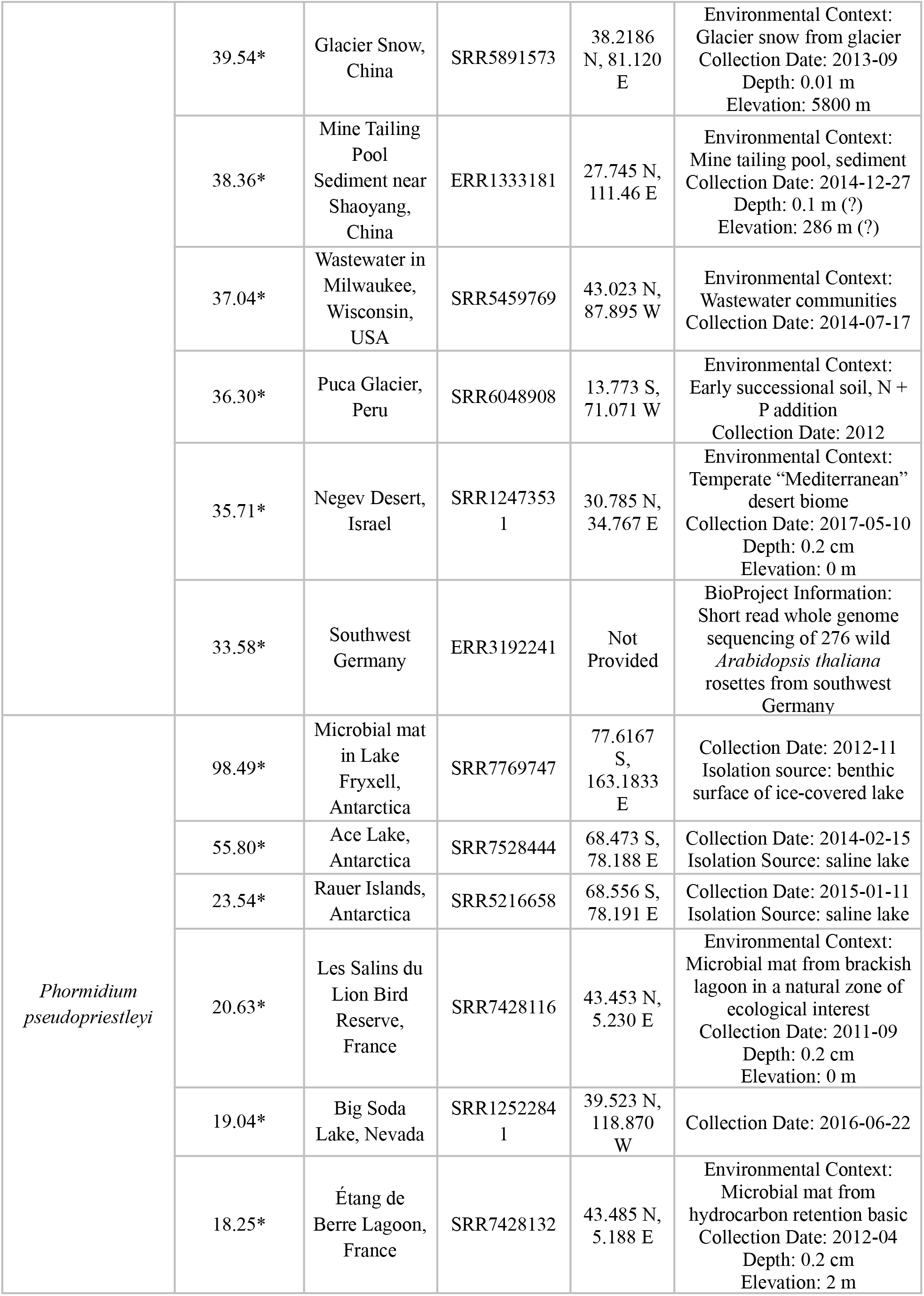
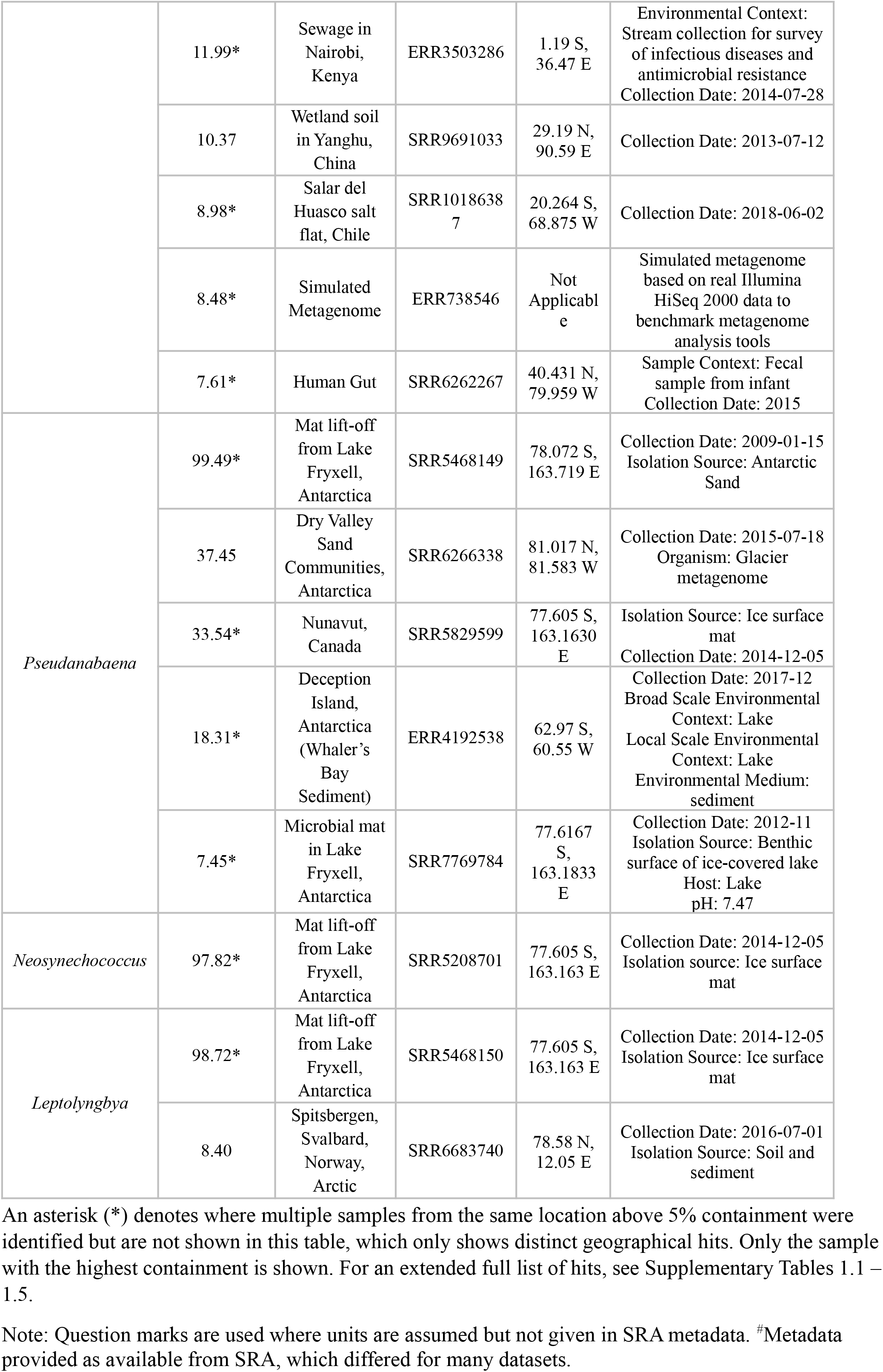
Metagenomes from the NCBI SRA with the Highest Containment Values

| MAG | Containment (%) | Location | Accession Number | Latitude and Longitude | BioSample Metadata<sup>#</sup> |
| --- | --- | --- | --- | --- | --- |
| *Microcoleus* | 99.18* | Mat lift-off from Lake Fryxell, Antarctica | SRR5468150 | 77.605 S, 163.1630 E | Isolation Source: Ice surface mat<br>Collection Date: 2014-12-05 |
|  | 65.02* | Polar Desert Sand Communities, Antarctica | SRR6266358 | 78.0741 S, 163.8918 E | Isolation Source: Antarctic Sand<br>Collection Date 2010-01-15 |
|  | 57.50* | Moab Green Butte, Utah, USA | SRR5855414 | 38.42 N, 109.41 W | Isolation Source: biocrust samples from Green Butte Site near Canyonlands National Park along an apparent maturity gradient of Cyanobacteria-dominated biocrusts<br>Collection Date: 2014-09<br>Sample Collection: soil coring with Petri dishes |
|  | 41.65* | Ningxia, China | SRR2952554 | Not provided | Isolation Source: algae crusts<br>Collection Date: 2013-04-15<br>Geographic Location: China: Ningxia |
|  | 41.10* | Sonoran Desert, Colorado Plateau, USA | SRR5247052 | 38.42 N, 109.4099 W | Isolation Source: Colorado Plateau and Sonoran Desert Plateau and Sonoran Desert |
|  | 40.61* | Pig Farm, UK | ERR3588763 | 55.95 N, -3.188 W | Environmental Context: Pig farm soil<br>Collection Date: 2017-01-11<br>Project Name: The dynamics of antimicrobial resistance gene prevalence on a commercial pig farm: implications for policy |
|  | 39.54* | Glacier Snow,<br>China | SRR5891573 | 38.2186<br>N, 81.120<br>E | Environmental Context:<br>Glacier snow from glacier<br>Collection Date: 2013-09<br>Depth: 0.01 m<br>Elevation: 5800 m |
|  | 38.36* | Mine Tailing<br>Pool<br>Sediment near<br>Shaoyang,<br>China | ERR1333181 | 27.745 N,<br>111.46 E | Environmental Context:<br>Mine tailing pool, sediment<br>Collection Date: 2014-12-27<br>Depth: 0.1 m (?)<br>Elevation: 286 m (?) |
|  | 37.04* | Wastewater in<br>Milwaukee,<br>Wisconsin,<br>USA | SRR5459769 | 43.023 N,<br>87.895 W | Environmental Context:<br>Wastewater communities<br>Collection Date: 2014-07-17 |
|  | 36.30* | Puca Glacier,<br>Peru | SRR6048908 | 13.773 S,<br>71.071 W | Environmental Context:<br>Early successional soil, N +<br>P addition<br>Collection Date: 2012 |
|  | 35.71* | Negev Desert,<br>Israel | SRR1247353<br>1 | 30.785 N,<br>34.767 E | Environmental Context:<br>Temperate "Mediterranean"<br>desert biome<br>Collection Date: 2017-05-10<br>Depth: 0.2 cm<br>Elevation: 0 m |
|  | 33.58* | Southwest<br>Germany | ERR3192241 | Not<br>Provided | BioProject Information:<br>Short read whole genome<br>sequencing of 276 wild<br>*Arabidopsis thaliana*<br>rosettes from southwest<br>Germany |
| *Phormidium*<br>*pseudopriestleyi* | 98.49* | Microbial mat<br>in Lake<br>Fryxell,<br>Antarctica | SRR7769747 | 77.6167<br>S,<br>163.1833<br>E | Collection Date: 2012-11<br>Isolation source: benthic<br>surface of ice-covered lake |
|  | 55.80* | Ace Lake,<br>Antarctica | SRR7528444 | 68.473 S,<br>78.188 E | Collection Date: 2014-02-15<br>Isolation Source: saline lake |
|  | 23.54* | Rauer Islands,<br>Antarctica | SRR5216658 | 68.556 S,<br>78.191 E | Collection Date: 2015-01-11<br>Isolation Source: saline lake |
|  | 20.63* | Les Salins du<br>Lion Bird<br>Reserve,<br>France | SRR7428116 | 43.453 N,<br>5.230 E | Environmental Context:<br>Microbial mat from brackish<br>lagoon in a natural zone of<br>ecological interest<br>Collection Date: 2011-09<br>Depth: 0.2 cm<br>Elevation: 0 m |
|  | 19.04* | Big Soda<br>Lake, Nevada | SRR1252284<br>1 | 39.523 N,<br>118.870<br>W | Collection Date: 2016-06-22 |
|  | 18.25* | Étang de<br>Berre Lagoon,<br>France | SRR7428132 | 43.485 N,<br>5.188 E | Environmental Context:<br>Microbial mat from<br>hydrocarbon retention basin<br>Collection Date: 2012-04<br>Depth: 0.2 cm<br>Elevation: 2 m |
|  | 11.99* | Sewage in Nairobi, Kenya | ERR3503286 | 1.19 S, 36.47 E | Environmental Context: Stream collection for survey of infectious diseases and antimicrobial resistance<br>Collection Date: 2014-07-28 |
|  | 10.37 | Wetland soil in Yanghu, China | SRR9691033 | 29.19 N, 90.59 E | Collection Date: 2013-07-12 |
|  | 8.98* | Salar del Huasco salt flat, Chile | SRR10186387 | 20.264 S, 68.875 W | Collection Date: 2018-06-02 |
|  | 8.48* | Simulated Metagenome | ERR738546 | Not Applicable | Simulated metagenome based on real Illumina HiSeq 2000 data to benchmark metagenome analysis tools |
|  | 7.61* | Human Gut | SRR6262267 | 40.431 N, 79.959 W | Sample Context: Fecal sample from infant<br>Collection Date: 2015 |
| *Pseudanabaena* | 99.49* | Mat lift-off from Lake Fryxell, Antarctica | SRR5468149 | 78.072 S, 163.719 E | Collection Date: 2009-01-15<br>Isolation Source: Antarctic Sand |
|  | 37.45 | Dry Valley Sand Communities, Antarctica | SRR6266338 | 81.017 N, 81.583 W | Collection Date: 2015-07-18<br>Organism: Glacier metagenome |
|  | 33.54* | Nunavut, Canada | SRR5829599 | 77.605 S, 163.1630 E | Isolation Source: Ice surface mat<br>Collection Date: 2014-12-05 |
|  | 18.31* | Deception Island, Antarctica (Whaler's Bay Sediment) | ERR4192538 | 62.97 S, 60.55 W | Collection Date: 2017-12<br>Broad Scale Environmental Context: Lake<br>Local Scale Environmental Context: Lake<br>Environmental Medium: sediment |
|  | 7.45* | Microbial mat in Lake Fryxell, Antarctica | SRR7769784 | 77.6167 S, 163.1833 E | Collection Date: 2012-11<br>Isolation Source: Benthic surface of ice-covered lake<br>Host: Lake<br>pH: 7.47 |
| *Neosynechococcus* | 97.82* | Mat lift-off from Lake Fryxell, Antarctica | SRR5208701 | 77.605 S, 163.163 E | Collection Date: 2014-12-05<br>Isolation source: Ice surface mat |
| *Leptolyngbya* | 98.72* | Mat lift-off from Lake Fryxell, Antarctica | SRR5468150 | 77.605 S, 163.163 E | Collection Date: 2014-12-05<br>Isolation Source: Ice surface mat |
|  | 8.40 | Spitsbergen, Svalbard, Norway, Arctic | SRR6683740 | 78.58 N, 12.05 E | Collection Date: 2016-07-01<br>Isolation Source: Soil and sediment |

An asterisk (\*) denotes where multiple samples from the same location above 5% containment were identified but are not shown in this table, which only shows distinct geographical hits. Only the sample with the highest containment is shown. For an extended full list of hits, see Supplementary Tables 1.1 – 1.5.
Note: Question marks are used where units are assumed but not given in SRA metadata. #Metadata provided as available from SRA, which differed for many datasets.

**Table 3.**
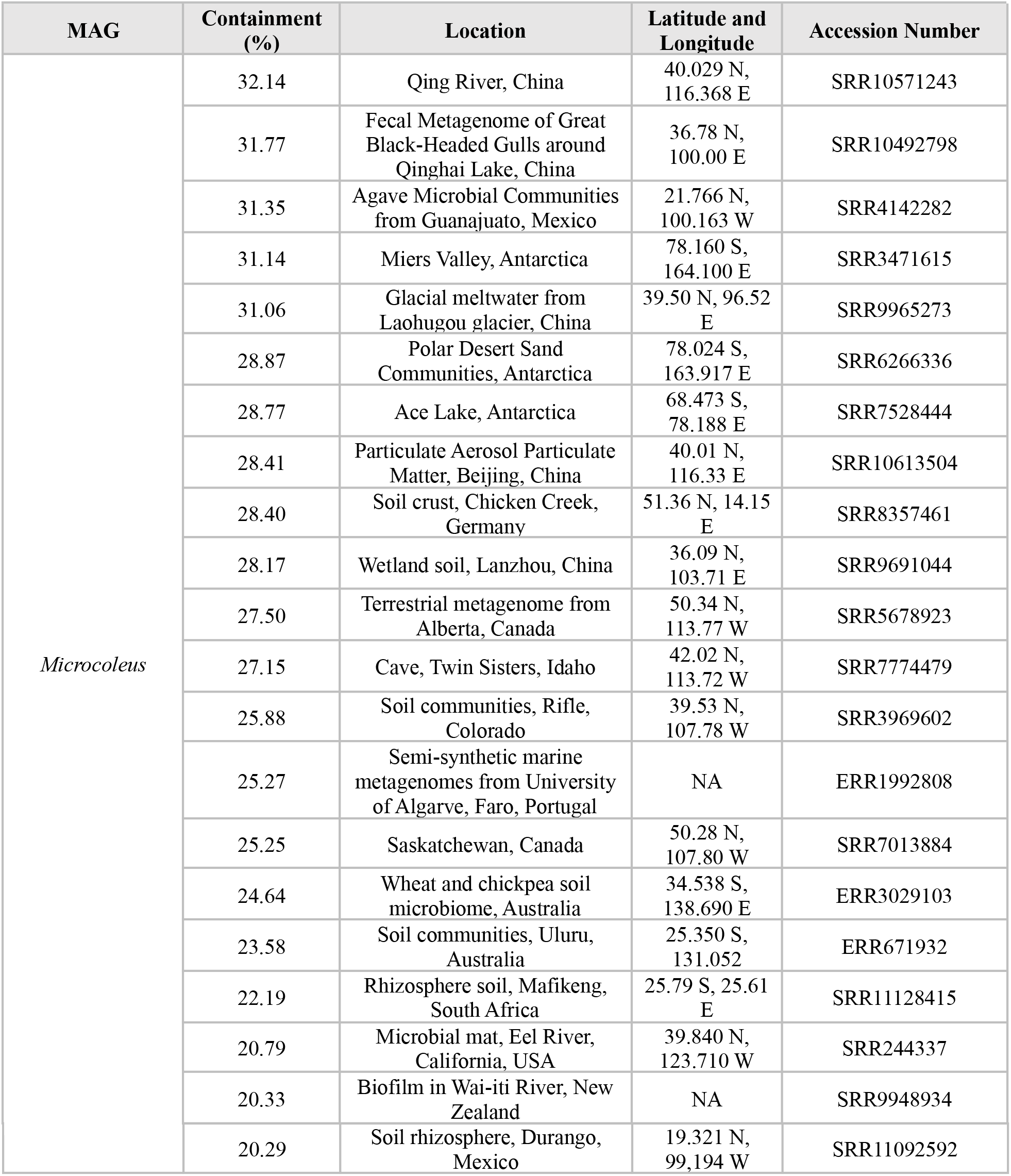
Additional Metagenomes from Unique Locations >20% Containment for *Microcoleus* MAG

**Table 4.**
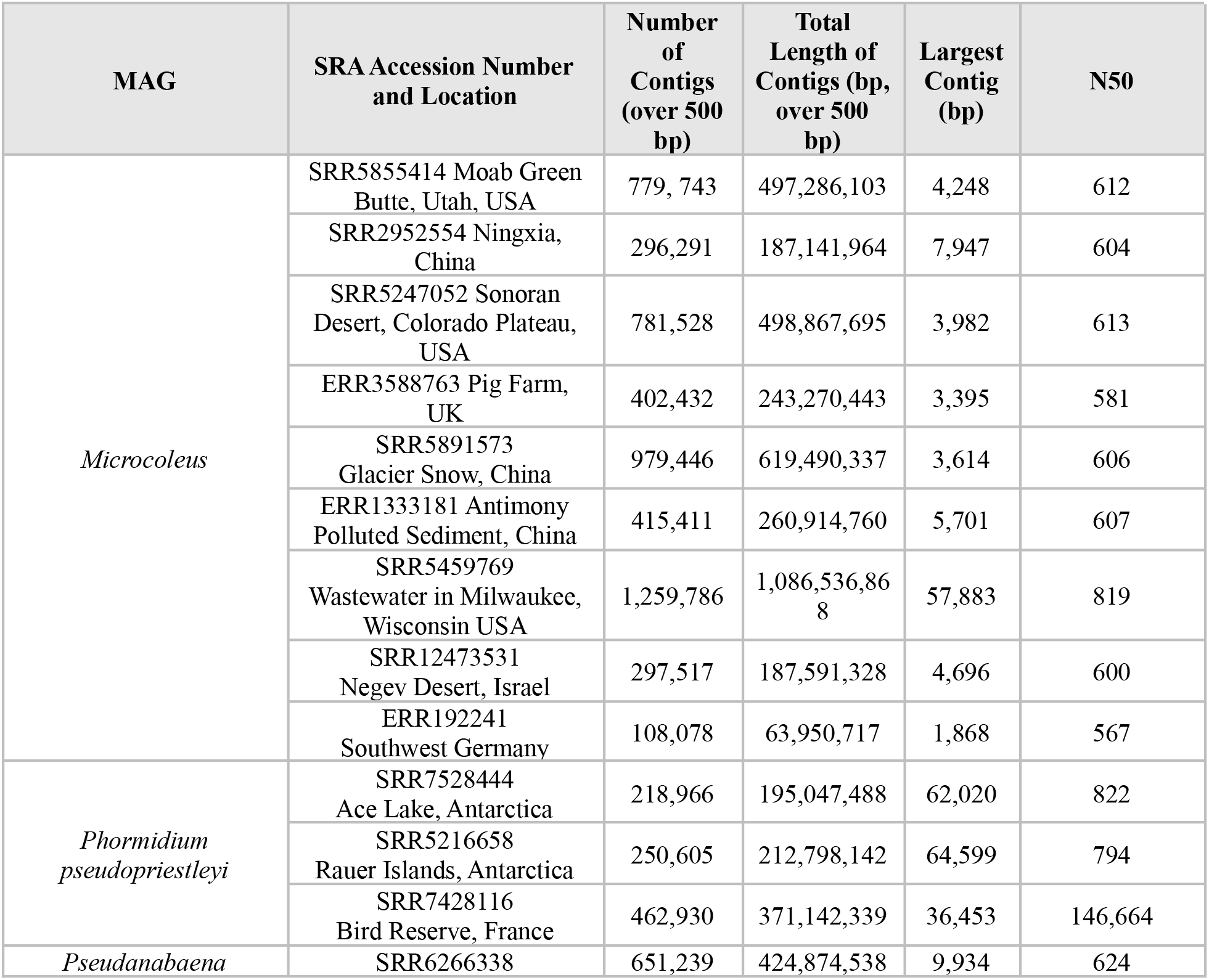
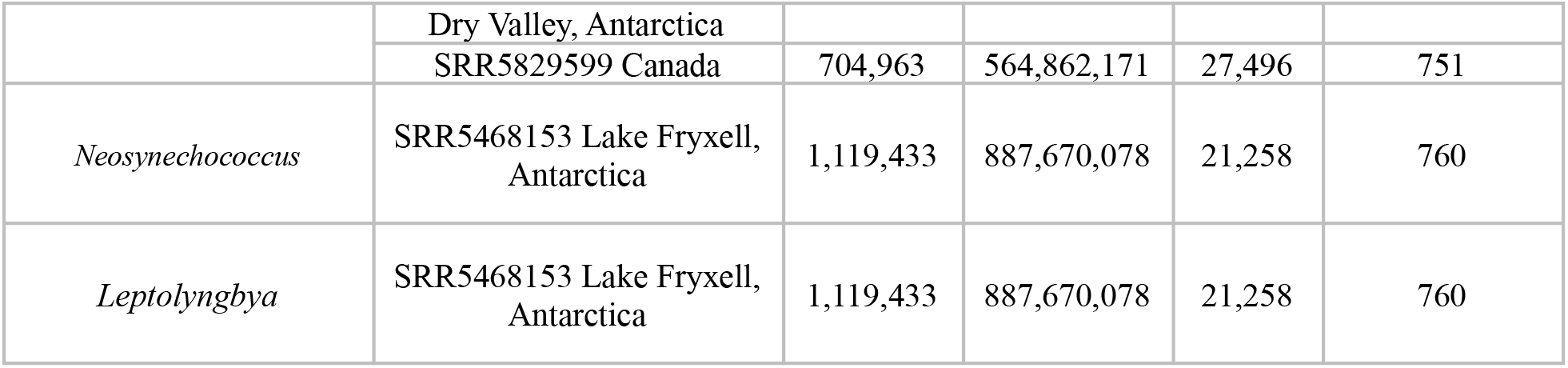
Quality Metrics of Metagenome Assemblies and Mapping Statistics

**Table 5.**
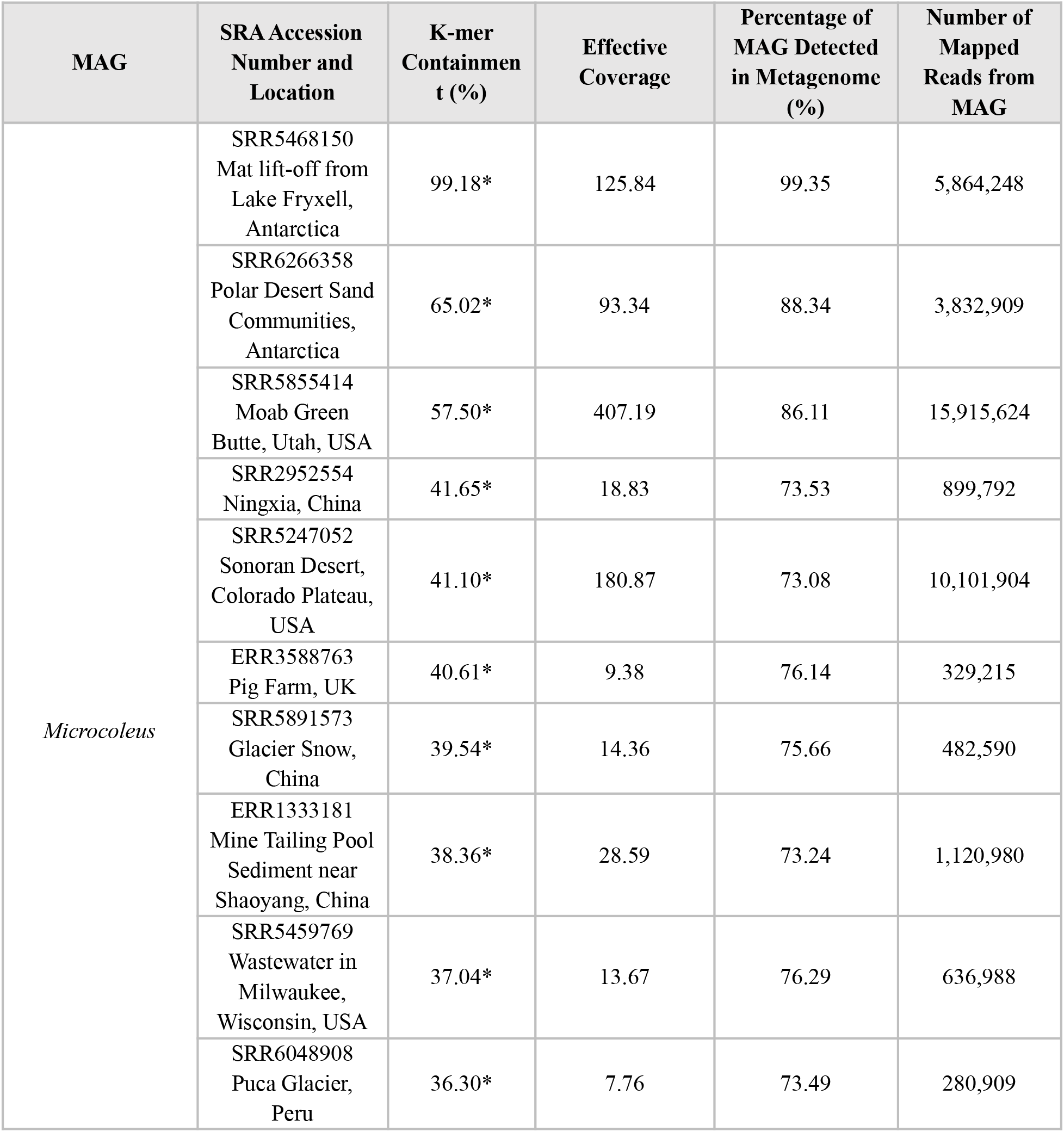
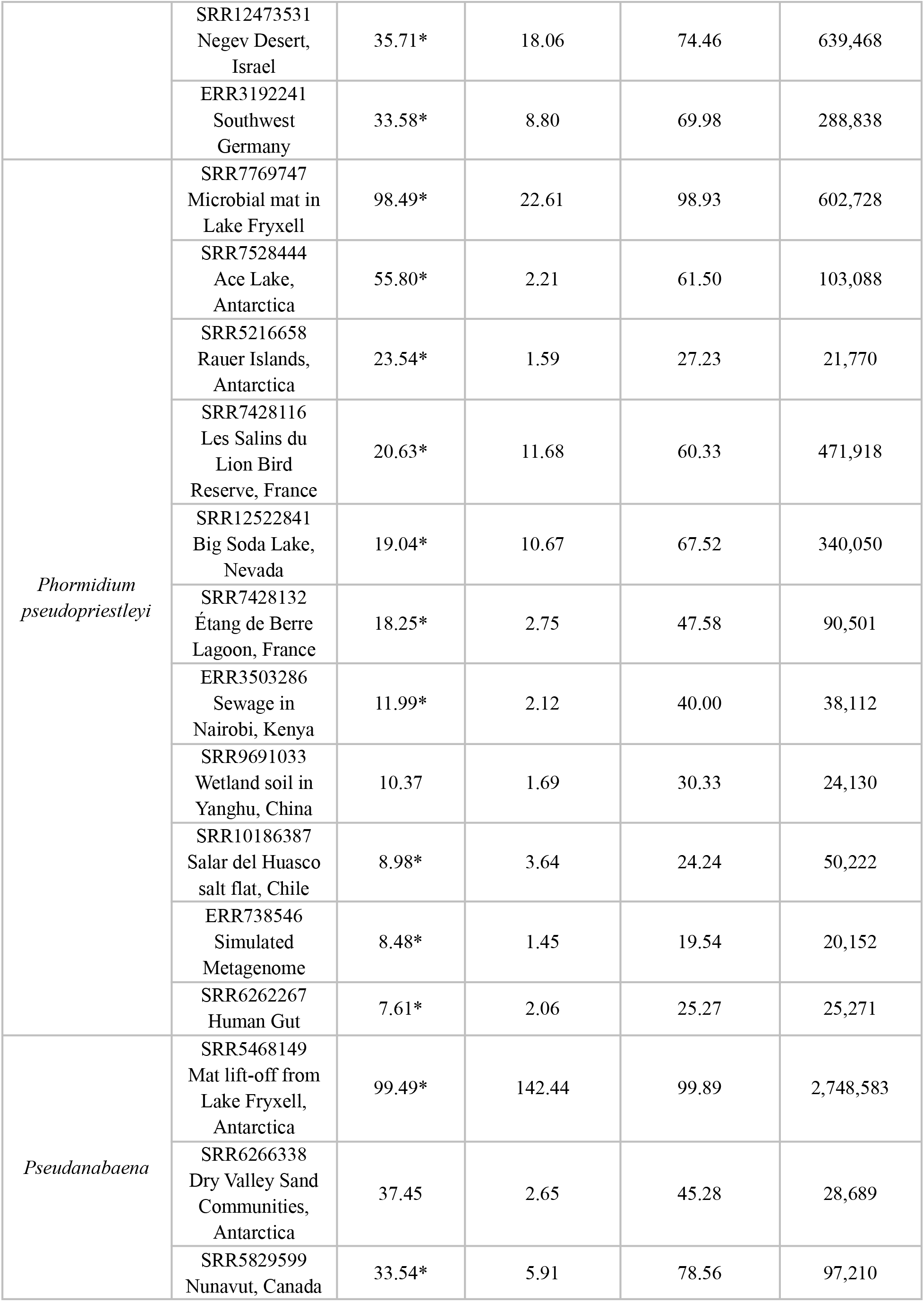
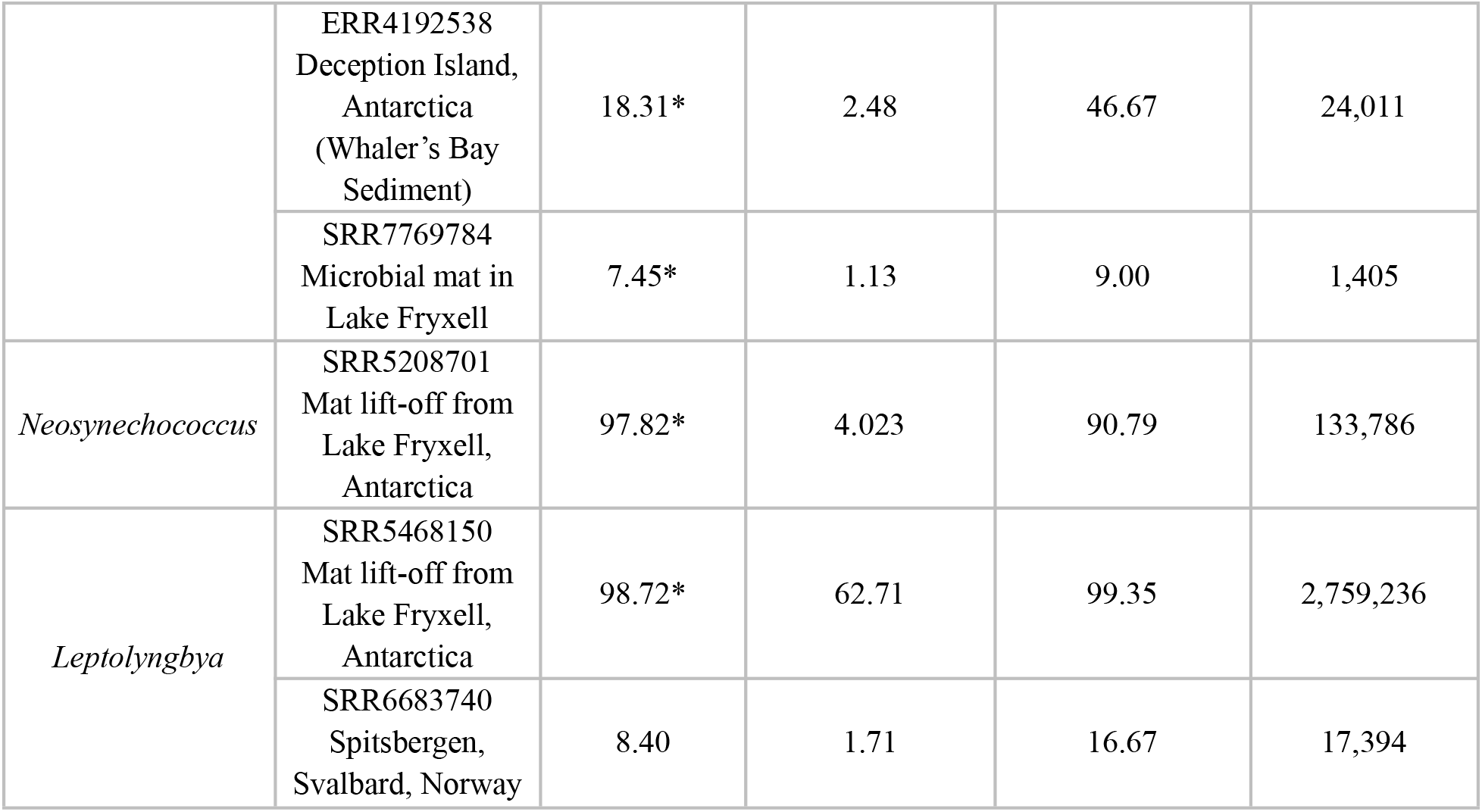
Mapping Validation of MAGs in SRA Metagenomes

**Figure 1.**
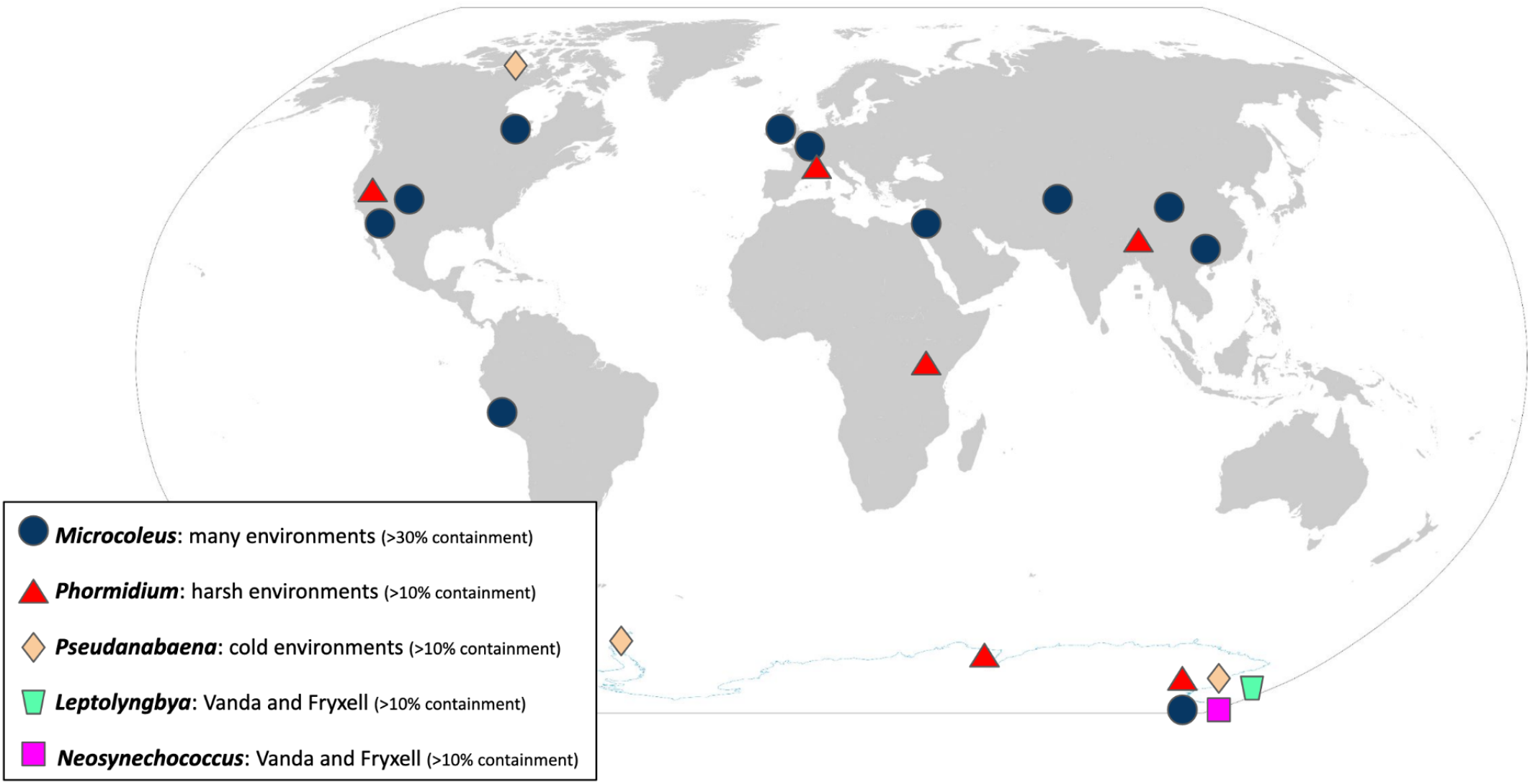
Global Distribution of Antarctic Cyanobacteria Global distribution of Antarctic MAGs above 30% for the *Microcoleus* MAG and above 10% for the remaining MAGs. The symbols over Lake Vanda and Fryxell are not positioned for scale to avoid crowding.

The purpose of branchwater was to find shared genomic data between Antarctic MAGs and SRA metagenomes from different habitats, geographic location, and climate zones. Matches of our selected Antarctic cyanobacteria MAGs in these metagenomes may indicate the occurrence of Antarctic cyanobacteria or closely related taxa in environments across the globe. A k-mer size of 31 with at least 5% containment indicates a ∼91% ANI between matched sequences (Hera et al., 2022). At 30% containment, this value increases to ∼97% ANI (Hera et al., 2022). Thus, a high containment value indicates the presence in the metagenome of genomic DNA similar to the MAG, and supports the presence of a related organism in the sampling location of that metagenome. Low k-mer containment values may represent smaller regions of shared genomic material or the presence of a related species, but cannot definitively support the presence of the same species in that environment. Containment, particularly at low values, can be affected by factors such as plasmids, metagenome sequencing depth, or small portions of shared contamination between the MAG and metagenome.

The *Microcoleus* MAG was the most widely distributed MAG with 27 globally distinct locations above 25% containment (Tables 1 – 3). The *Microcoleus* and *P. pseudopriestleyi* MAGs were present in the most time series and subsamples from the same environmental location, which resulted in 1,121 hits above 5% for the *Microcoleus* MAG and 131 hits for *P. pseudopriestleyi* MAG (Table 1). The *Pseudanabaena* and *P. pseudopriestleyi* MAGs were found in three distinct locations above 25% containment while the *Neosynechococcus* and *Leptolyngbya* MAGs were only found in one location above 25% containment (Table 1).

The *Microcoleus* MAG was found in diverse environments with conditions ranging from hot to cold climates and including both arid and wet locations (Tables 2 – 3). Some environments are cold year-round such as Puca Glacier in Peru (36.30% containment), glacier snow in China (39.54% containment), and the ice-covered Lake Vanda, while others are temperate, like Wisconsin, USA (37.04% containment), or Southwest Germany (33.58% containment). *P. pseudopriestleyi* MAG was found in three Antarctic metagenome data sets: Lake Fryxell mat samples (98.49% containment), Ace Lake (55.8% containment) and the Rauer Islands (23.54% containment). The highest 30 hits for the *P. pseudopriestleyi* MAG, including the three samples used to create the MAG, were from Lake Fryxell. This search revealed that *P. pseudopriestleyi* is likely present in other depths of Lake Fryxell than 9.8 m despite not being prevalent at those depths based on 16S sequencing (Jungblut et al., 2010). Besides Antarctica, the *P. pseudopriestleyi* MAG was found in a bird reserve next to a lagoon in France called Les Salins du Lion (20.63% containment) as well as a hydrocarbon polluted saline lagoon called Étang de Berre (18.25% containment) which were part of a study on the effects of hydrocarbon pollution on microbial communities (Aubé et al., 2016). The *P. pseudopriestleyi* MAG was also found in the Salar del Huasco salt flat in Chile (8.98% containment), antimicrobial treated sewage collected in Nairobi, Kenya (11.99% containment) and an infant gut fecal sample (7.61% containment). All these environments represent extreme conditions for cyanobacteria.

Although the *Microcoleus, Pseudanabaena, Neosynechococcus*, and *Leptolyngbya* MAGs were obtained from microbial mat pinnacles in Lake Vanda, they were all present in high containment (>97%) in mat lift-off samples from Lake Fryxell where the *P. pseudopriestleyi* MAG was not detected. The *Pseudanabaena* MAG was also found in a dry sand community in the McMurdo Dry Valleys (37.45% containment), where lakes Vanda and Fryxell are located, as well as Whaler’s Bay on Deception Island in Antarctic (18.31 % containment) and the Canadian High Arctic such as Nunavut, Canada (33.54 % containment), which is cold but geographically distant from the Antarctic.

Metagenomes representing geographically distinct locations were selected for further analysis to compare genomic data from different environments to the Antarctic MAGs. These data sets were run through an assembly and binning pipeline to obtain bins that could be compared to the Antarctic MAGs. However, metagenome assemblies were poor quality with the majority of the N50s under 1,000 base pairs, which is the minimum contig length required to bin with MetaBAT. Thus, bins were not generated, and it would not have been possible to identify the presence of the MAGs in these metagenomes without using an assembly-independent technique. Validation of the branchwater results was done by mapping the MAGs to metagenomes (Table 5). The percentage of the MAG detected in metagenome and average MAG coverage confirm the results of branchwater independent of k-mer comparisons, with all but one sample exhibiting higher mapping-based detection in the metagenome than k-mer containment.

## DISCUSSION

### Environmental Diversity of Microcoleus

The presence of the *Microcoleus* MAG in diverse environments indicates that it can survive in a range of different ecological conditions and climatic zones. The findings agree with previous biogeographic assessments of cultured cyanobacteria belonging to the species *Microcoleus vaginatus* and the *Microcoleus* spp. based on the 16S rRNA gene (Dvořák et al., 2012; StruneckÝ et al., 2013). In order to survive cold temperatures in Lake Vanda, the *Microcoleus* must deal with cellular membranes becoming brittle and slowed metabolism. However, some environments where the *Microcoleus* was found are only cold for part of the year (Moab Green Butte Desert; Ningxia, China; Southwest Germany; Milwaukee, Wisconsin; and the UK) while other environments are cold year-round (Puca Glacier, Peru, and glacial snow in China). In contrast to cold conditions, hot temperatures can cause proteins to denature and prolonged exposure to sunlight can cause high light and UV stress. These conditions occur in the Moab Green Butte Desert, the Sonoran Desert, and the Negev Desert. Furthermore, the Moab Desert and Sonoran Desert experience extreme temperature changes between morning and night (Turnage & Hinckley, 1938; Balling et al., 1998; McCann et al., 2018), forcing the *Microcoleus* to adapt to both conditions on a 24-hour cycle.

In addition to temperature range, the *Microcoleus* MAG was found in metagenomes environments with different levels of water availability and habitat types. Locations entailed arid desert soil crusts (Moab and Negev Deserts), mine tailings (Shaoyang, China; the United Kingdom; Milwaukee), epiphyte on plant microbiomes (Southwest, Germany), and freshwater environments (Qing River, China and Ace Lake, Antarctica). The *Microcoleus* MAG was also found in data from both high and low elevation environments (5800 m elevation in glacial snow in China and 0 m elevation in the Negev Desert). Interestingly, in Southwest Germany the MAG was found in metagenomic data of wild Arabidopsis plants. Overall, the variety of conditions where the *Microcoleus* MAG was found indicates that it may live in an impressive range of environments ranging from moderate climates to extreme heat or cold.

### Environmental Diversity of Phormidium pseudopriestleyi

*P. pseudopriestleyi* is a sulfide-tolerant cyanobacteria found in a low light environment in Lake Fryxell, Antarctica. Our study identified the *P. pseudopriestleyi* MAG in metagenomes from additional locations in Antarctica such as the saline Ace Lake (Vestfold Hills) and lakes on the Rauer Islands, which agrees with previous 16S rRNA gene sequencing where the species was documented from Salt Pond and Fresh Pond on McMurdo Ice Shelf (Jungblut et al., 2005; Lumian et al., 2021) as well as Ace lake (Taton et al., 2006). Interestingly, *P. pseudopriestleyi* or a close relative is present also at low abundance in a pond at Les Salins du Lion, a bird reserve (20.63% containment, 95% cANI), and Étang de Berre, a hydrocarbon polluted saline lagoon (18.25% containment, 94% cANI), both in southern France (Aubé et al., 2016). Four environmental conditions can be compared in these locations: irradiance, salinity, temperature, and sulfide concentrations. The irradiance at Les Salins du Lion pond and Étang de Berre lagoon was not measured when environmental sampling occurred, but the elevation of the lagoon was recorded to be at 0 m, indicating that irradiance is higher at the surface of the pond than the low irradiance at the depth of sampling in Lake Fryxell (1-2 µmol/photon m^−2^ s^−1^) (Sumner et al., 2015). Furthermore, Salt Pond and Fresh Pond have high illumination levels in the summer (Roos & Vincent, 1998; Jungblut et al., 2005), indicating that *P. pseudopriestleyi* may have the capability to overcome high irradiation and UV levels for prolonged periods. Les Salins du Lion (14 g L^−1^ NaCl) and Étang de Berre (20 g L^−1^ NaCl) have a lower salinity than Lake Fryxell (70.13 g L^−1^ NaCl) and Salt Pond (∼990 g L^−1^ NaCl), which is hypersaline (Jungblut et al., 2005; Aubé et al., 2016; Lumian et al., 2021). Previous work has showed that *P. pseudopriestleyi* increases the thickness of its extracellular polymeric substance layer in response to saline stress (Agrawal & Singh, 1999). Sulfide is also present in Les Salins du Lion, with a concentration of ∼0.24 g L^−1^ at the time of sampling (Aubé et al., 2016), which was the highest value at any location or time sampled included in the study. This demonstrates a much higher sulfide tolerance than what was previously recorded in the Lake Fryxell sampling site, which was 9.8 × 10^−5^ g L^−1^ (Lumian et al., 2021).

In addition to Les Salins du Lion and Étang de Berre, the *P. pseudopriestleyi* MAG was found in globally distributed challenging environments such as a salt flat in Chile, antimicrobial treated sewage in Kenya, and infant gut. The fact that *P. pseudopriestleyi* thrives in environments with harsh conditions suggests that it has capabilities to overcome diverse environmental stresses. In Lake Fryxell, *P. pseudopriestleyi* dominates microbial mats at 9.8 m depth in low light and sulfidic conditions but it is less abundant at shallower depths, even though there is more light availability and no sulfide (Jungblut et al., 2016; Dillon et al., 2020). Thus, *P. pseudopriestleyi* may grow slowly and find ecological success in environments that are too harsh for faster growing cyanobacteria, which is consistent with the slow growth rate of *P. pseudopriestleyi* seen in unpublished laboratory observations. The other environments where genomes similar to *P. pseudopriestleyi* were found may provide challenges that prohibit many other cyanobacteria from growing (polar environments, alkaline lake in Big Soda lake, antimicrobial treated sewage in Nairobi, Kenya), allowing *P. pseudopriestleyi* to survive in a nonpolar environment.

### Environmental Diversity of Pseudanabaena, Neosynechococcus, and Leptolyngbya

The top matches for the *Pseudanabaena, Neosynechococcus*, and *Leptolyngbya* MAGs showed that they were also present in Lake Fryxell and that the *Pseudanabaena* MAG was in sediment in the McMurdo Dry Valleys. The *Neosynechococcus* MAG was only present in the McMurdo Dry Valleys, however the presence of the *Leptolyngbya* and *Pseudanabaena* MAGs in geographically distant locations in the Arctic (Norway and Canada respectively) suggests that the cyanobacteria forming these MAGs have a global distribution in cold environments and might have undergone long range dispersal. The mechanism of long-range distribution could be wind; atmospheric studies show bacteria from the Saharan desert are transported by wind throughout the Atlantic (Griffin et al., 2002; Gorbushina et al., 2007; A. D. Jungblut et al., 2010). A similar process is expected to allow Antarctic cyanobacteria to cross large distances and populate diverse geographic regions. However, the lack of non-polar locations suggests that they are not as successful at integrating into non-polar environments. Thus, these cyanobacteria may be specific to polar environments even though they may be transported globally, which agrees with 16S rRNA gene analysis that proposed the presence of cosmopolitan cold ecotypes (Jungblut et al., 2010). The genus *Neosynechococcus* was described by Dvořák et al., (2014) based on cyanobacteria isolated from peat bog in Slovakia and was described based on 16S rRNA gene, 16S-23S ITS, and *rbcl* loci, however in the approached presented here it could only be identified in Antarctica.

### Implications for Biogeographic Distributions

The perceived distributions of organisms in biogeography studies are affected by sampling and publishing biases. Sampling in remote locations is logistically difficult and is often centered around established sampling locations which may be near research stations and infrastructure. This results in many studies and publications from established sampling locations and a deeper understanding of local ecology and geochemical processes in these environments. Biogeography studies, however, benefit from widespread sampling in many locations. Conducting widespread ecological sampling is expensive and can be impractical, so it is advantageous to search existing datasets for as much information as possible. Using branchwater to search public metagenomes makes the most out of data from remote areas by revealing previously unknown locations of organisms of interest. Furthermore, results from this analysis included remote areas, including various sites in Antarctica, which may not have otherwise been identified as locations of the query MAGs.

Despite being affected by sampling bias like all biogeography studies, the results showed that the *Microcoleus* MAG was globally distributed over a wide variety of environments, the *P. pseudopriestleyi* MAG was found in predominantly in harsh environments, the *Neosynechococcus* and *Leptolyngbya* MAGs were in the Antarctic, and the *Pseudanabaena* MAG was in geographically separated polar environments. The numerous sites containing the *Microcoleus* MAG imply that this species has the genetic capacity to adapt to many types of environments. It may also have a faster growth rate than an extreme conditions specialist, like *P. pseudopriestleyi* MAG, which would allow it to compete in a variety of ecological communities, some of which experience stressful conditions. Previous work has shown *Microcoleus* to be a cosmopolitan genus (Garcia-Pichel et al., 1996, 2001).

Although the *Microcoleus* MAG is by far the most globally diverse cyanobacterial genome in this study, there is variety in the distributions of the other four MAGs. The prevalence of the *P. pseudopriestleyi* MAG in harsh environments indicates that it finds ecological success in stressful environments, and it is likely outperformed by other organisms in moderate environments. The *Pseudanabaena, Neosynechococcus*, and *Leptolyngbya* MAGs were only found in polar environments, indicating they may be outcompeted in moderate environments. Diving deeper into the metabolic potential of each organism and interactions between metagenome community members may offer insights as to how and why some organisms are prevalent in a multitude of environments while others are prevalent in only certain conditions.

## CONCLUSIONS

This paper presents the first biogeography study using a large-scale k-mer-based approach and characterizes the global distribution of five distinct Antarctic cyanobacteria based on public data. We show that the *Microcoleus* MAG has cosmopolitan distribution and presence in a variety of environments, whereas the *P. pseudopriestleyi* MAG is also globally distributed but mostly present in harsh environments. *Leptolyngbya*, and *Pseudanabaena* MAGs were only found in polar environments from Arctic to Antarctica suggesting the existence of cosmopolitan cold ecotypes. The *Neosynechococcus* MAG was only detected in Antarctica and provides support for more restricted distribution patterns and potential endemicity. Further in situ transcriptomic studies of these MAGs may reveal adaptation mechanisms including why the *Microcoleus* is so pervasive compared to the other cyanobacteria in this study.

Branchwater can search ∼500,000 metagenomes with a query genome in under 24 hours on commodity hardware (Irber et al. unpublished). The ability to quickly find genomes similar to query MAGs in publicly available unassembled metagenomic data sets has important implications for biogeography studies, which have been predominantly based on 16S rRNA gene sequencing due to the prevalence of data and ease of comparison. Branchwater greatly increases the amount of data that can be used for biogeography studies. This technique is especially helpful for organisms that are in remote locations and underrepresented in genomic data, such as polar cyanobacteria, by providing a much larger number of known environments than would be possible with targeted field studies. Additionally, branchwater can be used to identify accessible sampling locations of organisms from remote environments, such as the *Microcoleus* being identified in the Moab Green Butte Desert in Colorado, USA at 41.10% containment. As more metagenome datasets are made publicly available on the NCBI SRA, more information about the distribution of cryosphere cyanobacteria can be attained. The results further demonstrate the potential of metagenomics and k-mer based MAG approaches in investigating biogeography and ecology of cyanobacteria and environmental microbiology in the Polar Regions.

