## Supplemental Tables 1-5 for "Biogeographic Distribution of Five Antarctic Cyanobacteria Using Large-Scale k-mer Searching with sourmash branchwater"

### SUPPLEMENT

### APPENDIX

**Supplemental Table S1.1** Extended Matches from branchwater *Neosynechococcus* MAG Hits >5% Containment

| MATCHES | CONTAINMENT | LOCATION |
|---|---|---|
| SRR5208701 | 97.82% | Lake Fryxell liftoff and glacier meltwater |
| SRR5468149 | 97.16% | Lake Fryxell liftoff and glacier meltwater |
| SRR5208700 | 81.58% | Lake Fryxell liftoff and glacier meltwater |
| SRR5208699 | 74.52% | Lake Fryxell liftoff and glacier meltwater |
| SRR5468150 | 65.46% | Lake Fryxell liftoff and glacier meltwater |
| SRR5468153 | 36.51% | Lake Fryxell liftoff and glacier meltwater |

**Supplemental Table S1.2** Extended Matches from branchwater *Leptolyngbya* MAG Hits >5% Containment

| MATCHES | CONTAINMENT | LOCATION |
|---|---|---|
| SRR5468150 | 98.72% | Lake Fryxell liftoff and glacier meltwater |
| SRR5468149 | 98.58% | Lake Fryxell liftoff and glacier meltwater |
| SRR5208699 | 98.35% | Lake Fryxell liftoff and glacier meltwater |
| SRR5208701 | 97.98% | Lake Fryxell liftoff and glacier meltwater |
| SRR5208700 | 97.70% | Lake Fryxell liftoff and glacier meltwater |
| SRR5468153 | 25.16% | Lake Fryxell liftoff and glacier meltwater |
| SRR6683740 | 8.40% | Arctic Lake Metagenome |

**Supplemental Table S1.3** Extended Matches from branchwater *Pseudanabaena* MAG Hits >5% Containment

| MATCHES | CONTAINMENT | LOCATION |
|---|---|---|
| SRR5468149 | 99.49% | Lake Fryxell liftoff and glacier meltwater |
| SRR5468150 | 99.46% | Lake Fryxell liftoff and glacier meltwater |
| SRR5208701 | 99.20% | Lake Fryxell liftoff and glacier meltwater |
| SRR5468153 | 99.13% | Lake Fryxell liftoff and glacier meltwater |
| SRR5208700 | 98.91% | Lake Fryxell liftoff and glacier meltwater |
| SRR5208699 | 98.12% | Lake Fryxell liftoff and glacier meltwater |
| SRR6266338 | 37.45% | Polar Desert Sand Communities |
| SRR5829599 | 33.54% | Nunavut, Canada |
| SRR5215118 | 30.57% | Nunavut, Canada |
| SRR5829597 | 26.16% | Nunavut, Canada |
| ERR4192538 | 18.31% | Deception Island, Antarctica (Whaler's Bay Sediment) |
| ERR4192539 | 16.43% | Deception Island, Antarctica (Whaler's Bay Sediment) |
| SRR7769784 | 7.45% | Antarctic Microbial Mat |
| SRR7769706 | 6.84% | Antarctic Microbial Mat |
| SRR7769810 | 5.46% | Antarctic Microbial Mat |

| | | |
|---|---|---|
| SRR8842248 | 5.39% | Cryoconite from Svalbard |
| SRR7769748 | 5.21% | Antarctic Microbial Mat |

**Supplemental Table S1.4** Extended Matches from branchwater *Phormidium* MAG Hits >5% Containment

| MATCHES | CONTAINMENT | LOCATION |
|---|---|---|
| SRR7769578 | 99.39% | Antarctic Microbial Mat |
| SRR7769621 | 99.32% | Antarctic Microbial Mat |
| SRR7769581 | 98.49% | Antarctic Microbial Mat |
| SRR7769747 | 98.49% | Antarctic Microbial Mat |
| SRR7769746 | 98.44% | Antarctic Microbial Mat |
| SRR7769635 | 98.25% | Antarctic Microbial Mat |
| SRR7769622 | 98.18% | Antarctic Microbial Mat |
| SRR7769634 | 98.11% | Antarctic Microbial Mat |
| SRR7769582 | 98.07% | Antarctic Microbial Mat |
| SRR7769583 | 97.79% | Antarctic Microbial Mat |
| SRR7769576 | 97.74% | Antarctic Microbial Mat |
| SRR7769579 | 97.60% | Antarctic Microbial Mat |
| SRR7769748 | 97.15% | Antarctic Microbial Mat |
| SRR7769793 | 96.56% | Antarctic Microbial Mat |
| SRR7769792 | 95.86% | Antarctic Microbial Mat |
| SRR7769639 | 94.70% | Antarctic Microbial Mat |
| SRR7769754 | 93.97% | Antarctic Microbial Mat |
| SRR7769518 | 92.65% | Antarctic Microbial Mat |
| SRR7769683 | 92.01% | Antarctic Microbial Mat |
| SRR7769753 | 91.90% | Antarctic Microbial Mat |
| SRR7769616 | 91.83% | Antarctic Microbial Mat |
| SRR7769558 | 91.00% | Antarctic Microbial Mat |
| SRR7769636 | 90.18% | Antarctic Microbial Mat |
| SRR7769554 | 88.88% | Antarctic Microbial Mat |
| SRR7769620 | 85.02% | Antarctic Microbial Mat |
| SRR7769624 | 83.72% | Antarctic Microbial Mat |
| SRR7769514 | 82.76% | Antarctic Microbial Mat |
| SRR7769557 | 81.55% | Antarctic Microbial Mat |
| SRR7769751 | 80.54% | Antarctic Microbial Mat |
| SRR7769794 | 77.45% | Antarctic Microbial Mat |
| SRR7769755 | 61.13% | Antarctic Microbial Mat |
| SRR7528444 | 55.79% | Ace Lake Saline |
| SRR7769788 | 54.53% | Antarctic Microbial Mat |
| SRR7529760 | 47.77% | Ace Lake Saline |
| SRR7769790 | 42.68% | Antarctic Microbial Mat |
| SRR7769643 | 39.13% | Antarctic Microbial Mat |
| SRR7769519 | 29.52% | Antarctic Microbial Mat |
| SRR7769812 | 25.39% | Antarctic Microbial Mat |
| SRR5216658 | 23.54% | Rauer Islands, Antarctica (saline) |
| SRR7769513 | 22.92% | Antarctic Microbial Mat |
| SRR6129595 | 22.38% | Rauer Islands, Antarctica (saline) |
| SRR7769618 | 22.10% | Antarctic Microbial Mat |
| SRR6185695 | 21.16% | Rauer Islands, Antarctica (saline) |
| SRR7529754 | 20.83% | Ace Lake Saline |
| SRR7428116 | 20.63% | Brackish Lagoon (SL) |
| SRR7428117 | 20.33% | Brackish Lagoon (SL) |
| SRR7428114 | 20.28% | Brackish Lagoon (SL) |
| SRR7769787 | 19.27% | Antarctic Microbial Mat |
| SRR7428121 | 19.17% | Brackish Lagoon (SL) |

| | | |
|---|---|---|
| SRR7428120 | 19.10% | Brackish Lagoon (SL) |
| SRR12522841 | 19.04% | Big Soda Lake, Nevada |
| SRR7428115 | 18.87% | Brackish Lagoon (SL) |
| SRR7769528 | 18.47% | Antarctic Microbial Mat |
| SRR7428132 | 18.25% | Brackish Lagoon (EBD) |
| SRR12522839 | 17.71% | Big Soda Lake, Nevada |
| SRR7769662 | 16.79% | Antarctic Microbial Mat |
| SRR7769657 | 16.75% | Antarctic Microbial Mat |
| SRR7769512 | 16.66% | Antarctic Microbial Mat |
| SRR12522840 | 16.52% | Big Soda Lake, Nevada |
| SRR7769693 | 16.11% | Antarctic Microbial Mat |
| SRR7529732 | 15.95% | Ace Lake Saline |
| SRR7769584 | 15.20% | Antarctic Microbial Mat |
| SRR7428131 | 14.98% | Brackish Lagoon (EBD) |
| SRR7769531 | 14.14% | Antarctic Microbial Mat |
| SRR7769802 | 14.11% | Antarctic Microbial Mat |
| SRR7769623 | 13.08% | Antarctic Microbial Mat |
| SRR7769574 | 13.01% | Antarctic Microbial Mat |
| SRR7769801 | 12.98% | Antarctic Microbial Mat |
| SRR7769678 | 12.70% | Antarctic Microbial Mat |
| SRR7769659 | 12.16% | Antarctic Microbial Mat |
| ERR3503286 | 11.99% | Nairobi, Kenya (sewage, antimicrobial resistance) |
| SRR7769811 | 11.76% | Antarctic Microbial Mat |
| SRR7529753 | 11.73% | Ace Lake Saline |
| SRR7769810 | 11.28% | Antarctic Microbial Mat |
| SRR7769559 | 11.26% | Antarctic Microbial Mat |
| SRR7769735 | 11.14% | Antarctic Microbial Mat |
| SRR7769804 | 10.96% | Antarctic Microbial Mat |
| SRR7769696 | 10.95% | Antarctic Microbial Mat |
| SRR7769515 | 10.89% | Antarctic Microbial Mat |
| SRR7769776 | 10.50% | Antarctic Microbial Mat |
| SRR9691033 | 10.37% | Yanghu, China (wetland soil) |
| SRR7769805 | 10.17% | Antarctic Microbial Mat |
| SRR7769660 | 10.11% | Antarctic Microbial Mat |
| SRR7769650 | 10.03% | Antarctic Microbial Mat |
| SRR7769752 | 10.03% | Antarctic Microbial Mat |
| SRR7769808 | 10.01% | Antarctic Microbial Mat |
| SRR7769734 | 9.47% | Antarctic Microbial Mat |
| SRR7769773 | 9.42% | Antarctic Microbial Mat |
| SRR7769575 | 9.07% | Antarctic Microbial Mat |
| SRR10186387 | 8.98% | Salar de Huasco, Chile (sediment) |
| ERR3503282 | 8.83% | Nairobi, Kenya (sewage, antimicrobial resistance) |
| SRR7769615 | 8.69% | Antarctic Microbial Mat |
| SRR7769580 | 8.64% | Antarctic Microbial Mat |
| SRR7769695 | 8.62% | Antarctic Microbial Mat |
| SRR7769775 | 8.51% | Antarctic Microbial Mat |
| SRR12522836 | 8.48% | Big Soda Lake, Nevada |
| ERR738546 | 8.48% | Simulated Metagenome |
| SRR7769553 | 8.41% | Antarctic Microbial Mat |
| SRR7769642 | 8.36% | Antarctic Microbial Mat |
| ERR738544 | 8.10% | Simulated Metagenome |
| ERR738545 | 7.87% | Simulated Metagenome |
| SRR7769664 | 7.87% | Antarctic Microbial Mat |
| SRR6262267 | 7.61% | Human Gut |
| SRR7428125 | 7.30% | Brackish Lagoon (EBD) |
| SRR7769736 | 7.05% | Antarctic Microbial Mat |
| SRR7769570 | 6.95% | Antarctic Microbial Mat |
| SRR7769803 | 6.48% | Antarctic Microbial Mat |

| | | |
|---|---|---|
| SRR11412982 | 6.31% | Human Gut |

**Supplemental Table S1.5** Extended Matches from branchwater *Microcoleus* MAG Hits >30% Containment

| MATCHES | CONTAINMENT | LOCATION |
|---|---|---|
| SRR5468150 | 99.18% | Lake Fryxell liftoff and glacier meltwater |
| SRR5468153 | 99.18% | Lake Fryxell liftoff and glacier meltwater |
| SRR5208700 | 98.66% | Lake Fryxell liftoff and glacier meltwater |
| SRR5468149 | 98.58% | Lake Fryxell liftoff and glacier meltwater |
| SRR5208699 | 86.19% | Lake Fryxell liftoff and glacier meltwater |
| SRR5208701 | 84.99% | Lake Fryxell liftoff and glacier meltwater |
| SRR6266358 | 65.02% | Polar Desert Sand Communities |
| SRR5855414 | 57.50% | Moab Soil Crust |
| SRR5855413 | 54.76% | Moab Soil Crust |
| SRR5855418 | 53.50% | Moab Soil Crust |
| SRR5855417 | 52.39% | Moab Soil Crust |
| SRR5855428 | 52.34% | Moab Soil Crust |
| SRR5855424 | 48.82% | Moab Soil Crust |
| SRR5855412 | 47.99% | Moab Soil Crust |
| SRR5855429 | 43.55% | Moab Soil Crust |
| SRR5855432 | 42.39% | Moab Soil Crust |
| SRR2952554 | 41.65% | Ningxia, China (soil crust) |
| SRR2954705 | 41.24% | Ningxia, China (soil crust) |
| SRR5247052 | 41.10% | Sonoran Desert |
| ERR3588763 | 40.61% | UK Pig Farm |
| SRR5855420 | 40.52% | Moab Soil Crust |
| SRR3439671 | 40.10% | Ningxia, China (soil crust) |
| SRR5830676 | 39.54% | Polar Desert Sand Communities |
| SRR5891573 | 39.54% | Glacial Snow, China |
| ERR1333181 | 38.36% | Mine Tailing Pool in China |
| SRR5459769 | 37.04% | Wastewater, Wisconsin |
| SRR6048908 | 36.30% | Puca Glacier, Peru |
| SRR12473531 | 35.71% | Mediterranean Desert Community |
| SRR12473532 | 35.68% | Mediterranean Desert Community |
| SRR5855438 | 34.78% | Moab Soil Crust |
| SRR12473534 | 34.30% | Mediterranean Desert Community |
| ERR3192241 | 33.57% | Southwest Germany (Arabidopsis community) |
